# Focal neuronal loss and cell replacement in a connectome model of fly steering

**DOI:** 10.64898/2026.10.26.762043

**Authors:** Pardes Habib, Michael Petr, Ruslan Rust

## Abstract

Focal neuronal loss after stroke, traumatic brain injury or spinal cord injury can disrupt circuits and result in persistent motor deficits. Which cells and connections must be restored to recover a specific behavior is difficult to establish experimentally. Here we use a connectome-constrained spiking model of the fruit-fly nervous system and decode motor-neuron output to simulate visually guided steering. After loss of 4,000 neurons, path straightness ranged from 0.04 to 0.95 depending on lesion location. At a fixed sensorimotor site, a 2,000-neuron lesion reduced simulated displacement to 6.5% of intact. Across 100 stochastic simulation realizations of the connectome, replacing 100 cells with their original connections increased displacement to 62.5% of intact when motor-control pathways were prioritized, compared with 10.1% after random selection of 100 cells and 48.5% after random selection of 1,000 cells. When connections were instead inferred from cell type, the same 100 intended replacements reached 60.0% of intact with matched identities and 6.7% with non-target identities, indicating that cell identity and type-dependent integration are required for recovery. Our model generates testable predictions for genetic ablation experiments in fruit flies and identifies candidate cellular and circuit requirements for motor repair that can inform cell replacement therapies and future models of mammalian locomotion.

## Introduction

Stroke and traumatic brain injury can cause focal neuronal loss and persistent circuit dysfunction^1,2^. The resulting deficits depend on which pathways are affected^3^. After stroke, for example, damage to the corticospinal tract predicts motor impairment beyond that explained by lesion volume alone^4^. Such observations help localize injury, but do not specify which neurons and connections would be needed to restore function. Transplanted neurons can receive host inputs and contribute to functional responses^5–7^, although their integration depends on the injured environment^8^. Defining the relative requirements for cell number, identity and connectivity is therefore central to developing and refining cell replacement therapies^9^.

The fruit fly, *Drosophila melanogaster*, offers an opportunity to model these requirements in a nervous system whose connections have been mapped at synaptic resolution. Recent reconstructions of the adult brain and complete male central nervous system (CNS) identify neurons and their connections, allowing lesions to be defined by the cells they remove^10–12^. The fly CNS contains the brain, optic lobes that process visual information, and a ventral nerve cord containing motor circuits. Descending neurons carry signals from the brain to these circuits^13,14^. This organization allows the effects of neuronal loss to be traced from sensory processing to motor output.

During flight, visual motion across the eyes provides information used to correct heading^15^. Direction-selective neurons, including T4 and T5 cells, detect this motion^16^. Downstream visual and descending pathways influence the motor neurons controlling the wings^13,14,17^. Models constrained by the fly’s synaptic connections can predict aspects of sensory and sensorimotor activity^18,19^. Visually guided steering is therefore a suitable task for testing how disruption and restoration of defined pathways affect movement.

Here we used a model of 165,122 fly neurons to predict how focal cell loss changes responses to visual motion. Motor-neuron activity determined turning and forward movement in a simplified simulation of steering. We varied lesion size and location, then tested whether a small population selected by circuit role restored more movement than a larger random population, and whether cell identity and connectivity mattered more than cell number. By comparing replacement strategies within the same injured circuit, we identify candidate requirements for cell therapy that can be tested by genetic ablation or silencing of defined cell types in adult flies and extended to more complex nervous systems.

## Results

### A circuit model connects visual motion to steering

We modeled 165,122 neurons and 124 million synapses from the male fly CNS connectome^12^. Each model neuron received signals through its mapped connections and generated electrical impulses, or spikes. Simulated visual motion activated direction-sensitive neurons, and wing motor activity determined steering (**Fig. 1a**). Responses to rightward and leftward motion were compared with forward motion to distinguish directional sensitivity from a persistent tendency to turn. A gain of one matched the intact calibration response; zero indicated no directional sensitivity.

**Figure 1:**
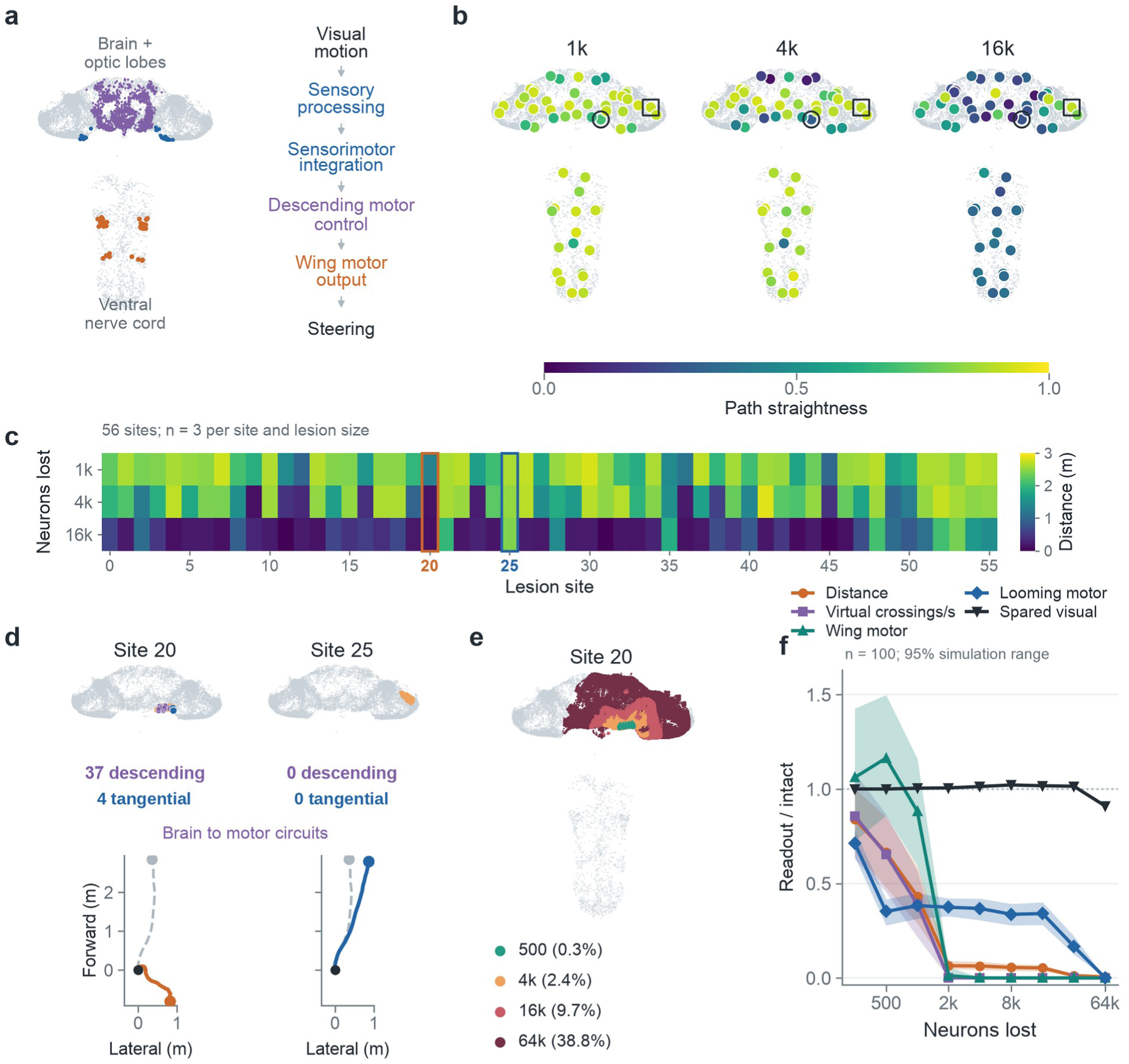
Lesion size and location determine movement impairment. **(a)** Soma positions and the sensorimotor pathway, shown at the same anatomical scale as b. **(b)** Mean path straightness after loss of 1,000, 4,000 or 16,000 neurons at 56 sites. Circles and squares mark sites 20 and 25. **(c)** Mean distance from the starting position after ten seconds at the same sites and lesion sizes. Columns follow numerical site order; outlines mark sites 20 and 25. Panels b,c average three neural seeds and 20 gusts per seed. **(d)** Matched 1,000-cell lesions and example paths. Purple marks lost descending neurons, blue lobula plate tangential cells and orange other lost cells. Dashed paths show intact. Examples use neural seed 1000 and gust seed 3. **(e)** Nested site-20 lesions; percentages refer to all 165,122 model neurons. **(f)** Mean readouts with the central 95% simulation range, defined by the 2.5th to 97.5th percentiles across 100 realizations. Movement averages 20 gusts before percentile calculation. Readouts are normalized to matched intact. Black shows the spared visual response. Wing-motor activity uses forward-motion input; looming activity measures the takeoff pathway. Virtual crossings count passages in either direction through four flight planes. These are neural and controller readouts; wing posture and observed takeoff are not simulated. Absolute units appear in **Supplementary Fig. 3a–d**. Residual movement at zero neural drive reflects the controller’s minimum speed.

Motor responses determined turning and forward movement in a two-dimensional simulation exposed to matched wind disturbances. We measured distance from the starting position after ten seconds and path straightness, the distance between the start and endpoint divided by total path length. Straightness is one for a straight route and approaches zero when movement makes little net progress. In 100 intact realizations, distance was 2.59 ± 0.17 m (mean ± s.d.; **Supplementary Fig. 1a**), and mean straightness was 0.916. Changing the gust sequence altered the path of the same intact circuit (**Supplementary Fig. 1b,c**). These trajectories describe steering without full flight biomechanics. Lesion and replacement outcomes are expressed relative to the matched intact network unless absolute units are given.

To assess agreement with experimental observations, we tested model counterparts of three published fly visual-pathway perturbations (**Supplementary Fig. 2a–c**). Silencing the three right HS neurons reduced rightward steering output by 20.6%, consistent with impaired optomotor yaw responses after unilateral HS lesions in blowflies^20^; T4/T5 silencing abolished steering output, as expected because these cells receive the simulated motion input, in agreement with the loss of optomotor turning reported in Drosophila^21^. LPLC2 silencing reduced giant-fiber firing by 19.8%, consistent with its contribution to looming-evoked giant-fiber responses^22^.

### Lesion location and size determine the modeled deficit

A screen compared focal loss of 1,000, 4,000 and 16,000 neurons at 56 fixed locations. Lesions removed neighboring cell bodies while preserving motor neurons as the output population. Median straightness across locations decreased from 0.905 to 0.880 and 0.345 as lesion size increased (**Fig. 1b**). After loss of 4,000 neurons, mean straightness ranged from 0.042 to 0.952 across sites. Distance from the starting position decreased between 1,000- and 16,000-cell lesions at 55 locations (**Fig. 1c**); at central-brain site 35, distance was 55% of intact after 1,000 lost neurons and 71% after 16,000. Thus, lesion size influenced overall impairment, while location influenced the deficit at each size.

We selected posterior central-brain site 20, an impaired site whose lesion removes both descending neurons and lobula plate tangential cells, to examine focal loss of descending motor-control pathways. In flies, visual descending neurons combine tangential-cell outputs and project directly to flight motor centers^23^. The 1,000-cell lesion at site 20 removed 37 descending neurons and four lobula plate tangential cells (**Fig. 1d**). The lost descending neurons included the giant fiber (DNp01) of the escape takeoff pathway^24^. Descending neurons carry brain signals to motor circuits in the ventral nerve cord. After this lesion, leftward gain was 0.16 and rightward gain was 1.01. A 1,000-cell lesion at optic-lobe site 25 spared both groups and retained 99.8% of intact distance, compared with 43.2% at site 20 (**Supplementary Movie 1**).

Selective removal of the 37 descending neurons retained 57.4% of intact distance, compared with 101.0% after removing 37 other cells sampled from the same lesion mask. Removing only the four tangential cells retained 98.0% (**Supplementary Fig. 3e,f**). These cell-class ablations help distinguish contributions within the focal lesion; they do not assume that every descending neuron specifically controls flight.

Across the nested site-20 series, distance fell from 84.0% of intact after 250 lost neurons to 6.5% after 2,000 and 0.58% after 64,000 (**Fig. 1e,f** and **Supplementary Movie 1**). The readouts differed in their size dependence. After 500 cells were removed, wing-motor activity was 116.4% of intact, despite distance falling to 66.3%. This decline followed the looming-evoked takeoff output, which contributes to motor drive and fell to 35.1%. The candidate spared visual response retained 101.4% of intact after 32,000 lost neurons and 91.0% after 64,000. Its relative preservation provides a specificity control in **Fig. 1f** and **Supplementary Fig. 3d**. Absolute distances, crossing rates and neural rates are shown in **Supplementary Fig. 3a–d**. At 64,000 lost neurons, wing-motor and takeoff outputs were zero; residual displacement came from the controller’s minimum speed.

### Prioritizing lost cells improves repair with original connections

We compared random selection, projection priority and motor-control priority when choosing which lost cells to restore (**Supplementary Fig. 4a**). Projection priority selected descending neurons and cells carrying signals into or out of the optic lobes. Motor-control priority first selected descending neurons and lobula plate tangential cells, followed by other optic-lobe projection cells. Each selected cell received its original identity and connections to available partners. This comparison identifies useful cells under an ideal repair rule. **Supplementary Movie 2** shows looming-evoked spikes and motor activity for one prespecified realization.

After a 2,000-cell lesion, replacing 100 cells selected by motor-control priority increased distance to 62.5% of intact (95% confidence interval 60.9–64.0%), compared with 10.1% after random selection (9.1–11.1%) and 20.3% after projection priority (18.4–22.1%; **Supplementary Fig. 4a** and **Supplementary Movie 3**). The selected 100-cell population comprised 79 descending neurons, five tangential cells and 16 other visual neurons, and therefore included every descending neuron lost in the lesion.

For the 2,000-cell lesion, original-wiring replacement ranged from 50 to 2,000 cells, with larger grafts retaining the cells present at smaller doses (**Supplementary Fig. 4a**). The 100-cell selected graft produced 1.61 m of displacement, compared with 1.25 m after 1,000 randomly selected replacements. At 2,000 cells, the three original-wiring selection rules share complete reconstruction of the lost population. Complete original reconstruction reproduced intact output exactly (**Supplementary Fig. 1d**). This endpoint defines an ideal repair benchmark. Intermediate doses are simulated outcomes; no plateau or monotonic response was imposed. In an earlier exploratory grid, 2,000 replacements with random nearby partners after a 1,000-cell lesion produced 0.30 m of displacement across three realizations (**Supplementary Fig. 4d**). This excess dose duplicates cell templates and changes synaptic load, unlike complete reconstruction.

### Replacement identity and connections shape functional recovery

We next compared original repair, random wiring, correct-type grafts and non-target grafts at the same 100 intended replacement positions, alongside no graft (**Fig. 2a–c** and **Supplementary Movie 4**). Original repair reached 62.5% of intact distance, whereas random nearby partners reached 8.7% (8.0–9.4%). The cell-therapy comparison assigned either the required identities or non-target central-brain identities, using the same type-based integration rule, which assumes that grafted neurons connect like endogenous neurons of their type^25^. Non-target identities came from annotated central-brain interneuron types absent from the lesion, excluding sensory, visual-projection, descending and motor classes. Copying partners from surviving neurons of the matched types reached 60.0% (58.3–61.6%), compared with 6.7% (6.3–7.0%) with non-target types. On average, 89 of the 100 matched cells copied their partners from an opposite-side neuron of the same type, with partners reflected, so for most cells this rule approximates the original wiring through bilateral symmetry. Assigned identity changed the resulting connections and synaptic load, testing identity together with type-dependent integration.

**Figure 2:**
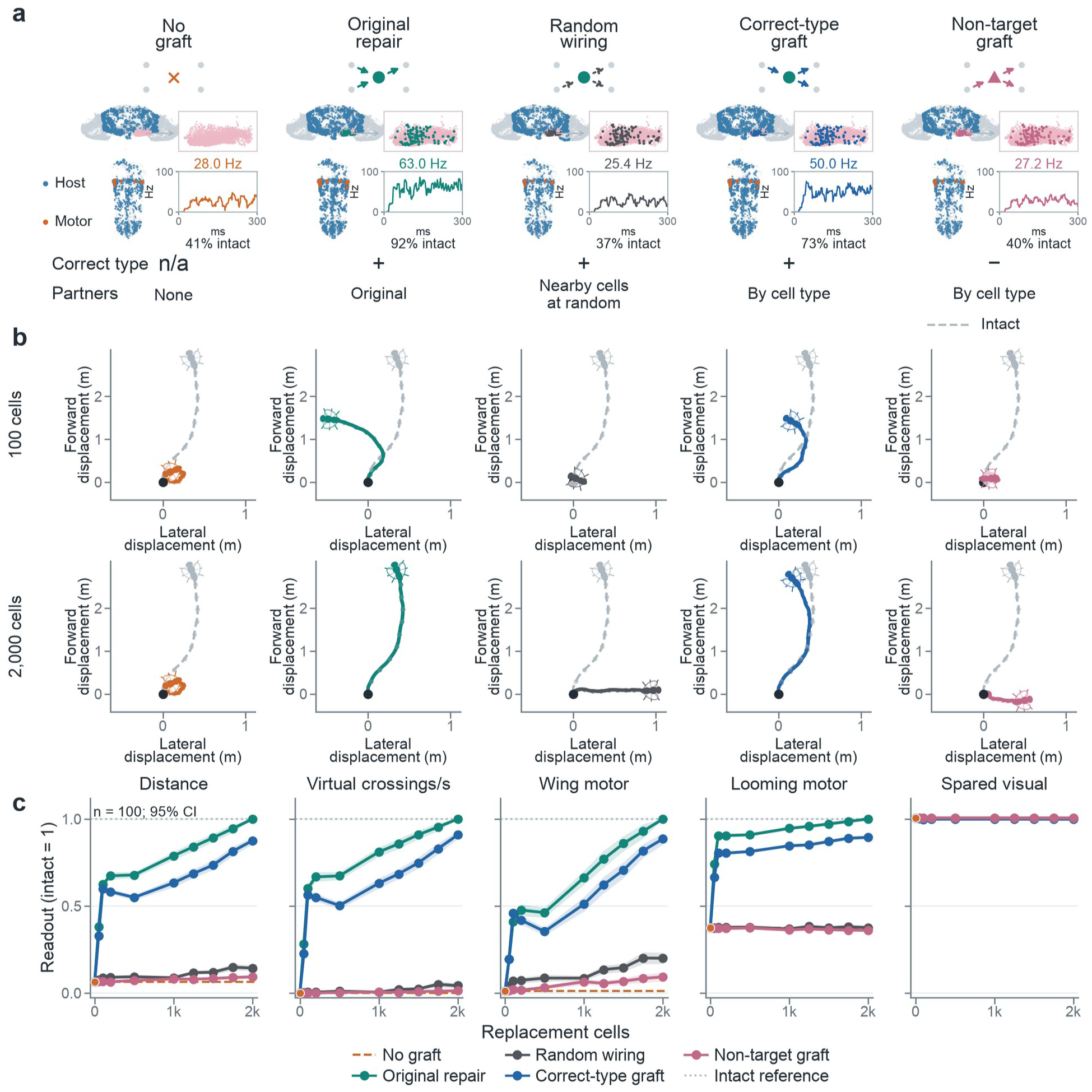
Cell identity and synaptic wiring determine replacement benefit. **(a)** Circles denote matched identities, triangles non-target identities and the cross no graft. Original repair restores available partners; random wiring samples nearby cells. “By cell type” takes partners from a surviving cell of the assigned type. Pink marks the 2,000-cell lesion; grafts share 100 intended replacement positions. Brain and ventral nerve cord share scale. Blue and orange show host and takeoff-motor looming activity (50–300 ms) for one matched realization. Numbers give mean motor rates in Hz and % intact. Boxes show trailing 25-ms motor-rate traces; marker size increases with rate. Anatomy does not depict wiring differences. **(b)** Ten-second paths with 100 or 2,000 replacements, using neural seed 1000, intervention seed 2000 and gust seed 3. Gray dashed paths show intact. Fly size is constant and posture illustrative; dots mark seconds. Paths share origins and axis ranges. **(c)** Readouts relative to matched intact. Zero dose shows damage (orange dashed); intact is one (gray dotted). Points and ribbons show means and pointwise 95% confidence intervals across 100 realizations, averaging 20 matched gusts first. Lines join tested doses without smoothing. At full dose, 42 untyped positions receive sampled lost-cell identities in the correct-type graft.

The four repair rules were tested from 50 to 2,000 replacements, with the same intended positions at each dose (**Fig. 2b,c**). At 2,000 cells, displacement reached 87.6% with matched identities and 9.3% with non-target identities. Random wiring reached 14.3% (95% confidence interval 12.7– 16.0%), whereas complete original repair reproduced intact output. Thus, replacing as many cells as were lost was insufficient when their identities or partners differed from those of the original circuit. The correct-type graft used sampled lost-cell identities at the 42 untyped positions. An earlier control that shuffled lost identities between positions is retained in **Supplementary Fig. 4c**. These grafts still supply useful missing types and reached 38.0% of intact at 1,000 and 52.9% at 2,000 replacements.

The dose curves start at the measured function remaining after damage, with matched intact defined as one (**Fig. 2c**). Distance, virtual crossings and wing-motor activity were markedly reduced, whereas the looming response retained 37.4% of intact activity. Movement recovery did not imply complete restoration of neural responses. With original connections, the selected 100-cell graft restored wing-motor activity to 40.9% of matched intact, while the looming-evoked takeoff-pathway response reached 90.5% (**Fig. 2c**). By contrast, the spared-side visual response was 100.6% after the untreated lesion (100.5–100.6%). Its relative preservation alongside a large motor deficit supports response specificity within the tested stimuli.

Directional responses and the shuffled-lost-type control provide additional tests of integration (**Supplementary Fig. 4b,c**). After the 2,000-cell lesion, leftward responses exceeded intact in all five 100-cell conditions, so the best grafts improved movement without correcting this steering imbalance. Earlier exploratory tests with three realizations examined original-partner retention, replacement survival and variable synaptic strength (**Supplementary Fig. 4d–f**). Together, these controls suggest that dose, cell survival, identity and integration each contribute to replacement benefit.

## Discussion

Focal injuries leave different repair requirements because they remove different cells and interrupt different pathways. This is relevant to stroke, traumatic brain injury and spinal cord injury, where neuronal loss and disconnection can produce persistent motor deficits^1,2,26^. In the fly model, lesion size predicted the overall decline in movement, while location determined which steering pathways were lost. Replacing a small, selected population restored more movement than replacing a larger, randomly selected population, and replacing as many cells as were lost was insufficient when their identities or connections differed. These findings suggest that cell number should be considered together with the circuit roles of the cells being supplied.

Several features of the simulated deficits are qualitatively consistent with earlier experiments in flies. Blocking the T4 and T5 motion detectors abolishes optomotor turning^16,21^, and losing the horizontal lobula plate tangential cells, through mutation or unilateral microsurgical lesion, reduces optomotor yaw responses^20,27^. The four tangential cells removed here are a small part of this population, which may explain why their loss had little effect. In walking flies, manipulating a single descending neuron type on one side can shift steering toward that side^28^. In the model, removing only the 37 descending neurons left directional gains close to intact, so other cells in the site-20 lesion contribute to its direction-specific loss. Silencing other descending neurons whose activity predicts turning leaves turning intact^29^, suggesting redundancy within the descending population, and during flight the DNg02 descending neurons regulate wingbeat amplitude as a population^30^. We are not aware of an in silico lesion of a fly connectome model whose behavioral predictions have been tested experimentally. In *C. elegans*, connectome-based predictions of neurons required for locomotion were confirmed by single-cell ablation^31^. The three targeted comparisons qualitatively reproduce selected perturbation effects on fly steering and looming-pathway activity (**Supplementary Fig. 2**).

Experimental transplantation studies support several parts of this interpretation. Grafted neurons can receive sensory inputs and contribute to motor behavior after stroke^6,7^. In spinal cord injury, the regional identity of neural grafts influences corticospinal regeneration and functional integration^26^. Yet cell therapy can also act through effects beyond neuronal replacement, including trophic and immunomodulatory bystander effects in the brain and in peripheral immune organs^1,32^. Human neural progenitor grafts in stroke-injured mice have been associated with improved movement, vascular repair and changes in inflammatory and graft-host signaling^33^. Our simulations isolate the contribution of added neurons and their connections. They do not capture these other therapeutic effects.

Improved behavior does not necessarily indicate restored circuit function. Replacing 100 selected neurons improved the path substantially, although wing-motor output and directional responses remained abnormal. A more normal trajectory therefore did not establish restoration of the original circuit. In experimental stroke, recovery can involve recruitment of new corticospinal connections after growth-promoting treatment and training, or enhanced cortical sprouting after delivery of GDF10^34,35^. These findings illustrate why movement should be assessed alongside circuit activity and connectivity. Our model measures the immediate consequences of specified replacement rules; it does not simulate axonal regeneration, rehabilitation or the biological processes through which compensation develops.

Incomplete replacement nevertheless produced benefit under several connection rules. Benefit depended on cell identity and the integration rule. Shuffled lost identities still helped at larger doses because they supply missing types, whereas non-target identities did not. Experimental evidence that delayed transplantation can improve early graft survival and proliferation after stroke adds a biological constraint that the present model does not predict^36^. Survival, identity, maturation and integration should therefore be measured separately and in relation to the timing of transplantation^37^.

Clinical studies emphasize the same need to distinguish improvement from its mechanism. The single-arm PISCES-2 study examined intracerebral neural stem-cell implantation after stroke but could not establish efficacy against an untreated control^38^. In chronic traumatic brain injury, the randomized STEMTRA trial reported improvement in motor impairment after modified mesenchymal stromal-cell implantation; its 48-week report included post hoc analyses and did not establish neuronal circuit replacement as the mechanism^39^. Our findings may help formulate circuit-level hypotheses for such research, but they do not specify a clinical cell product, dose or treatment window.

Several assumptions limit the quantitative predictions. All neurons share simplified electrical properties, whereas detailed models of human and rat cortical pyramidal cells show that dendritic structure and nonlinear synaptic integration alter how inputs become spikes^40^. Matching replacement-cell physiology may therefore matter in addition to identity and connectivity. The model also depends on its operating point: reducing synaptic amplitude by 15% abolished intact wing-motor output during forward stimulation, making normalized movement undefined at that setting. The collapse of wing-motor output after larger lesions may therefore partly reflect this proximity to threshold. Motor drive also combines wing-motor output with looming-evoked takeoff output, so loss of the giant-fiber pathway reduces simulated forward speed; this coupling is a simplification of the controller. When motor drive was based on wing-motor output alone or held constant, selected cells still outperformed random selection and matched identities still outperformed non-target identities. These comparisons therefore held across controller mappings, although their magnitude varied. Somatic removal omits axonal injury, inflammation, vascular changes and glial responses^41^. In the fly, cell bodies lie in an outer layer around the neuropil, so a lesion defined by the nearest somata does not map onto a single contiguous region of synaptic tissue, as a vascular infarct would. Motor neurons are protected, and repeated simulations sample variability within one reconstructed animal. The steering simulation lacks ongoing neural visual feedback and full flight biomechanics; coupling to a more detailed body model would test these assumptions^42^.

Experimental tests could combine visual-evoked left-right wingbeat asymmetry, flight distance and heading with recordings of motor and visual-pathway activity. Virtual crossing rates and tracked flight distance provide candidate behavioral endpoints, while looming-evoked motor activity can be compared with recordings from the takeoff pathway. Video-based tracking could provide continuous measures of these endpoints after targeted neuronal manipulation, as used for deep learning-based profiling of stroke recovery in rodents^43^. Predicting their absolute values, latency or response probability would require a stimulus-onset model, which is not included here. In the fly, these predictions could also be tested genetically. The class-selective lesions modeled here, such as loss of the 37 descending neurons compared with 37 other cells from the same region, could be reproduced by ablating defined cell types in adult flies. Candidate targets include the giant fiber and the other posterior descending neurons lost at site 20. Reversible silencing that is later released returns the same cells with their original connections, approximating the original-connection repair rule. Continuous monitoring over the following days could then separate the immediate deficit predicted here from later compensation, which the model does not simulate. Existing fly injury models, such as hypoxia-reoxygenation, damage the whole animal^44^; genetic ablation would more closely match the focal lesions simulated here. Mammalian connectomes provide detailed local cortical circuits^45,46^, but modeling locomotion will also require the relevant sensory, descending, spinal and motor pathways. These simulations define candidate cell populations and connections for testing motor repair after a specified injury.

## Methods

### Connectome and neuronal activity

We used the MaleCNS v1.0 connectome^12^, retaining 165,122 traced neurons, 25,563,197 directed connections and 124,025,046 anatomical synapses. Cell identities and annotations were retained, and soma coordinates were converted from 8-nm voxels to micrometers. Predicted GABA, glutamate and histamine outputs were assigned inhibitory signs; other or missing transmitter predictions were assigned excitatory signs^47^. Synapse counts and transmitter predictions approximate connection strength and sign.

Each neuron accumulated synaptic input and generated spikes using a leaky integrate-and-fire model based on a published fly network model^18^. Membrane potential relative to rest, v, and signed synaptic input, g, followed dv/dt = (−v + g)/20 ms and dg/dt = −g/5 ms. Crossing 7 mV reset both states to zero and imposed a 2.2-ms refractory period, during which states were held and arriving inputs discarded. Directly stimulated cells had no refractory period. The timestep was 0.1 ms and synaptic delay 1.8 ms, with analytic integration between events. Every 1 ms, cells outside refractoriness with both state magnitudes at most 0.001 mV were set to zero.

Each anatomical synapse contributed 0.14390733 mV. This value scaled the source model’s 0.275-mV amplitude by the ratio of average synapse counts per neuron in the source and MaleCNS networks, without fitting lesion outcomes. External input had a separate amplitude of 68.75 mV. The compiled simulator was checked against a NumPy reference for spikes and final states.

### Sensory input and motor responses

We tested each network with five separate 300-ms stimulus trials, measuring firing rates from 50 to 300 ms. Rightward rotation activated T4/T5 a subtypes on the right and b subtypes on the left; leftward rotation used the mirrored populations. Forward translation activated bilateral a subtypes. Separate trials activated gustatory cells or LC4/LPLC2 cells that respond to an approaching object (looming). Each input slot received an independent event with probability 0.01 per timestep, approximating 100-Hz Poisson stimulation. Lost inputs continued to consume random draws so that surviving cells received matched event sequences. Correctly assigned one-to-one sensory replacements inherited the original input slots.

Turning commands were calculated from the 32 wing-steering motor neurons. We took their mean right-minus-left response across five intact simulations, antisymmetrized it across matched left/right cell types, and normalized the resulting weights to unit length. The weighted sum of motor-neuron rates gave steering command y. These weights were fixed for all lesions and replacements. Directional responses were y_R_ − y_F_ and y_F_ − y_L_ for rightward and leftward motion, respectively, where F denotes forward motion. Dividing by the corresponding intact mean response gave gains G_R_ and G_L_. Turning bias was B = 2(y_F_ − y_F,0_)/(y_R,0_ − y_L,0_), where subscript 0 denotes the intact reference. Both intact calibration spans were positive.

Wing-motor output was the mean firing rate of left and right power-motor groups during forward stimulation, divided by the intact reference. A separate takeoff-related output summed the TTMn and DLMn group-mean rates during looming stimulation and was normalized in the same way. Their mean defined motor drive. Evoked activity counted neurons with measured spikes outside optic-intrinsic and optic-sensory classes, normalized each stimulus to intact and averaged the five values. Replacement cells contributed to their assigned populations, using the original population sizes as denominators. These readouts have no experimentally validated threshold for flight or survival.

**Supplementary Movie 2** shows the late part of the looming-stimulus spike recordings for the intact model, the 2,000-cell lesion and 100 replacements with original connections. The nine-second excerpt covers 59–300 ms of the 300-ms trial. All use the first prespecified realization, neural seed 1000 and intervention seed 2000. Traces show the arithmetic mean of TTMn and DLMn population rates over the preceding 25 ms, consistent with the absolute looming motor readout. The normalized wing-motor measure instead uses forward-motion input. Left tangential-cell activity during leftward input, LPTC_L_L, was specified as a candidate spared response before the main simulations and divided by its matched intact rate. **Figure 2a** maps host and takeoff-motor firing in the same 50–300-ms looming window for one matched realization of each 100-cell repair condition, plotting only known soma positions. Numbers average TTMn and DLMn population rates, relative to matched intact. Profiles use a trailing 25-ms window, as in **Supplementary Movie 2**.

### Comparison with published perturbations

Comparisons with published perturbations (**Supplementary Fig. 2**) used the unchanged neural parameters and steering decoder, with 10 matched sensory-event seeds (1000–1009) per intact/perturbed pair. Silencing targeted the three right HS neurons (HSN, HSE and HSS), all 13,580 T4/T5 neurons, or all 185 LPLC2 neurons. Stimuli were 300-ms trials with 100-Hz Poisson input; responses were measured over 50–300 ms. Rightward T4/T5 input was used for panels a,b and combined LC4/LPLC2 input for panel c. These additional simulations were separate from the 7,900 neural/intervention realizations.

### Lesion size and location

We tested 56 fixed spatial centers. At each center, the nearest 1,000, 4,000 or 16,000 eligible somata were silenced, producing nested lesions. Of 140,024 neurons with soma coordinates, 139,218 were eligible after motor neurons were excluded. All 815 annotated motor neurons were protected as a common output population. Neurons without coordinates remained in the network but could not be removed.

Site 20 was selected as vulnerable during earlier exploratory work and held fixed for detailed tests. The nine lesion sizes were 250, 500, 1,000, 2,000, 4,000, 8,000, 16,000, 32,000 and 64,000 neurons. Site 25 was the predetermined location control. The exploratory 56-site screen used three neural seeds per condition. The main simulations used 100 realizations for the site-20 size series, the three site-25 examples, replacement selection and connection tests, intact controls and complete reconstruction. Within the 1,000-cell site-20 mask, separate ablations removed all 37 descending neurons, all four tangential cells, their union, or 37 or four other cells sampled without replacement. These class-selective controls match cell counts but are not lesions of matched spatial volume. Cell annotations and direct synaptic targets are supplied in the source tables.

### Choosing replacement cells

New model neurons were added to the surviving network and assigned to cells lost in the lesion. Random selection sampled these lost cells without replacement. Projection priority selected descending, visual-projection and visual-centrifugal neurons. Motor-control priority selected lobula plate tangential cells and descending neurons, followed by other visual-projection and visual-centrifugal cells, then the remaining population. Order within each tier was randomized. For the 2,000-cell lesion, the first 100 cells under this rule comprised 79 descending neurons, five tangential cells and 16 other visual neurons.

Partial replacement restored 50, 100, 200, 500, 1,000, 1,250, 1,500 or 1,750 cells after the 2,000-cell lesion. Within each realization and selection rule, one permutation per priority tier supplied nested prefixes across all doses. The 2,000-cell endpoint was evaluated once as complete original reconstruction and shared across the three selection curves. Connection and type-assignment comparisons used exactly the same 100 templates chosen by motor-control priority within each realization.

A typed-dose series tested matched identities and shuffled lost identities at 50, 200, 500, 1,000, 1,250, 1,500, 1,750 and 2,000 cells, reusing the 100-cell conditions. Intended positions followed the same ranked prefixes. Identity and donor draws remained dose-specific; the resulting typed connections were not nested. Of the 2,000 lost cells, 1,958 had annotated types and 42 were untyped. At an untyped position, the algorithm sampled an annotated lost-cell type in either arm. Identity matching therefore refers to positions with known types; the first 100 intended positions were all annotated.

A further series used non-target central-brain types at all nine doses, including 100 cells. Eligible types were represented exclusively by central-brain intrinsic neurons and absent from the annotated lesion population. Sampling uniformly with replacement from 23,997 eligible surviving cells weighted the 5,809 types by their abundance. The intended positions and donor-based wiring operations matched the correct-type comparison. No outcome-based filtering of this pool or imposed performance ceiling was used.

A random-wiring series added eight dose conditions: 50, 200, 500, 1,000, 1,250, 1,500, 1,750 and 2,000 cells, reusing the original 100-cell condition. Intended identities and motor-control-priority positions were retained while synaptic partners were sampled locally. Together, the original 46 conditions and the 16 typed-dose, nine non-target and eight random-wiring additions comprise 79 conditions, 7,900 neural/intervention realizations and 158,000 nested movement trajectories.

Earlier three-seed controls separately varied complete-dose connection rules and, after a 1,000-cell lesion, combined 500, 1,000 or 2,000 replacement cells with original-partner probabilities of 0, 0.25, 0.5, 0.75 or 1. Those exploratory excess-dose results are presented separately.

### Connecting replacement cells

Under the original-connection rule, each connection retained its original partner with probability p. Otherwise, it was assigned to a surviving neuron within twice the lesion radius. Connections to lost cells without a replacement were omitted. Copied connections retained their synapse counts and presynaptic signs, and connections between replacement cells were generated once from their source. Replacing each lost cell exactly once with all original partners recreated the intact network, providing an implementation check. Doses exceeding the number lost duplicated cell templates and linked all corresponding internal pairs, so total synaptic load was not fixed across the dose-by-partner experiment.

For connections inferred from cell type, each replacement received either its intended identity or an identity sampled from the specified pool. The non-target pool contained only central-brain types absent from the annotated lesion. Earlier shuffled-type controls instead sampled other lost types, weighted by their abundance. Partners were copied from a surviving neuron of the assigned type, preferentially on the same side. For an opposite-side donor, partners were mapped to corresponding reflected cells where available. If no donor was available, local partners were sampled using the lost cell’s synapse budgets. Assigned type determined output sign and membership in the measured populations. Conflicting incoming and outgoing proposals for a connection between replacement cells were averaged, and self-connections were omitted. Only correctly assigned sensory cells inherited the original sensory input.

Survival tests initially added 100 or 200 selected cells, with each retained independently at probability 0.5 or 0.75. We recorded the actual number surviving. Separate tests retained 100 or 2,000 cells but varied their synaptic strengths. Each cell received a multiplier exp(σz − σ²/2), with z drawn from a standard normal distribution and σ equal to 0.5 or 1. The expected per-cell multiplier was one, but total graft strength was not fixed. Connections between replacement cells used the geometric mean of the two cell multipliers.

### Movement simulation and analysis

The measured gains, bias and motor drive determined movement in a two-dimensional steering controller for ten seconds. Angular velocity ω produced command −3G_dir_ ω_sat_ tanh(ω/ω_sat_), with ω_sat_ = 300 degrees per second. Positive angular velocity used G_L_ and negative angular velocity used G_R_. Bias added 150B degrees per second. Wind disturbances had a 150-ms correlation time and baseline amplitude of 100 degrees per second. Angular velocity relaxed over 20 ms and was integrated at 1-ms steps. Motor drive, clipped to 0.05–1, multiplied a nominal forward speed of 0.3 m per second and inversely scaled disturbance amplitude. This minimum speed permits residual displacement even when neural motor drive is zero. Neural responses were measured once per realization; the controller used these fixed gains in closed loop, without re-simulating the network or modeling flight biomechanics. **Supplementary Movies 1, 3 and 4** replay these saved trajectories at 60 frames per second from a fixed camera above a 0.5-m grid, with positions interpolated from the saved paths without smoothing. The fly is enlarged; its posture, altitude and wingbeat are illustrative.

For the separate controller checks in **Supplementary Fig. 1b,c**, we reused the 100 intact neural states while setting gust amplitude to 0, 0.5, 1, 1.5 or 2 times its standard value. Each state used the same 20 gust seeds, giving 10,000 diagnostic trajectories, including 2,000 repeated standard-amplitude trajectories. This exploratory analysis added no neural states and is separate from the 158,000 trajectories in the replacement and lesion batch. Standard amplitude reproduced the existing intact outcomes. The no-gust example retained the circuit’s existing gains, bias and drive.

Distance from the starting position was the Euclidean separation between the first and last saved positions. Straightness was this distance divided by total path length. Each realization was averaged across 20 matched gust sequences. Normalized measures divided each realization’s outcome by its matched intact outcome; absolute distances remain in meters. Values above intact were retained. Virtual gates were placed at 0.5, 1.0, 1.5 and 2.0 m along the initial forward axis, perpendicular to that axis. A gate counted once if any saved trajectory point reached or exceeded its position. This unique-gate count ranges from zero to four. A separate descriptive crossing-rate endpoint counted repeated passages in either direction through the same planes. A passage required movement from at least 5 mm on one side to at least 5 mm on the other, suppressing repeated counts near a boundary. Total passages were divided by the ten-second window. Crossing rate can increase with recrossing and is not a pure measure of forward progress. The absolute looming-evoked motor rate was the arithmetic mean of the TTMn and DLMn population rates. No stimulus-off baseline was simulated, so this is an evoked neural response rather than a baseline-subtracted locomotor effect.

The main simulations used neural seeds 1000–1099 and separate intervention seeds 2000–2099. Conditions were paired by realization and gust sequence; neural randomness varied sensory events, while intervention randomness varied cell order, donor choice or partner choice. Repeating an identical specification reproduces the same result. Decoder weights and intact population calibration remained fixed from the original five calibration seeds, 0–4. The independent unit for uncertainty was the neural/intervention realization, not each of its 20 trajectories. Except for **Fig. 1f** and the individual-outcome displays in **Supplementary Fig. 4e,f**, uncertainty displays show means and pointwise 95% confidence intervals, calculated as the mean plus or minus the 97.5th percentile of the Student t distribution multiplied by the standard error across independent realizations. **Figure 1f** instead shows means with the central 95% simulation range, defined by the empirical 2.5th and 97.5th percentiles of the 100 realization outcomes. Movement is averaged over 20 gusts before calculating these percentiles. These ribbons describe outcome variability rather than precision of the mean. Paired effects were calculated within realization before their confidence intervals. Source tables also report the 2.5th and 97.5th percentiles of realization outcomes. These intervals describe stochastic variability conditional on one connectome and fixed model parameters, not biological replication or model validity. No outcome-dependent exclusions or curve smoothing were applied.

Separate sensitivity tests changed synaptic amplitude by ±15% or input rate by ±25%, using the original neural seed 0 and a matched intact reference at each setting (archived sensitivity source tables). When intact wing-motor output was zero, normalized movement was undefined. Directional-response similarity was the mean of C(G_R_) and C(G_L_), with C(x) = max[0, 1 − |x − 1|], penalizing deficient and excessive gain. These operating-point tests were not pooled with the main simulations. A separate controller sensitivity analysis replayed the stored gains, bias and motor outputs of each realization through the same controller and 20 gust sequences, with motor drive defined by wing-motor output alone or held constant (controller sensitivity source table). The survival and strength analyses in **Supplementary Fig. 4e,f** retain three original seeds and show every outcome with its mean, without confidence intervals.

## Data and code availability

The MaleCNS v1.0 connectome is available from Janelia FlyEM. Model inputs derived from it, all simulation outputs and per-realization readouts, figure and movie source data, and the code used for simulations, analyses, figures and movies are available at https://github.com/rustlab1/fly-connectome-cell-replacement.

## Author contributions

R.R. designed the study and wrote the code. P.H. contributed to the experimental design and writing. M.P. contributed to the experimental design and writing. All authors agreed to and approved the final manuscript.

