## Supplementary Information for "Focal neuronal loss and cell replacement in a connectome model of fly steering"

Ruslan Rust, Ph.D.

Zilkha Neurogenetic Institute, Keck School of Medicine

University of Southern California

1501 San Pablo Street, Los Angeles, CA 90033, USA

**ORCID**

Pardes Habib: 0000-0002-5771-216X

Michael Petr: 0000-0002-1808-0163

Ruslan Rust: 0000-0003-3376-3453

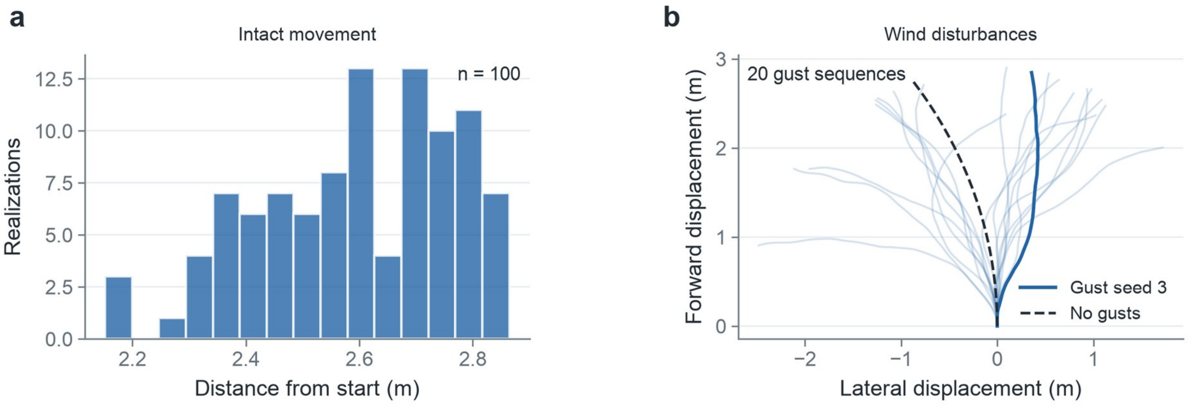


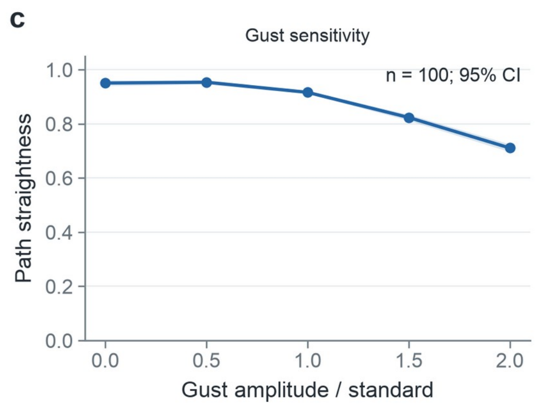


**d**

Original reconstruction

| Readout | Maximum difference |
|---|---|
| Steering commands | 0 |
| Wing motor | 0 |
| Looming motor | 0 |
| Distance | 0 |

Compared with matched intact; n = 100

**Supplementary Figure 1: Controller and reconstruction checks. (a)** Intact distance after ten seconds across 100 neural realizations, each averaged over 20 gusts. **(b)** Twenty gust sequences applied to the same intact neural realization (seed 1000), with gust seed 3 highlighted and a no-gust reference. **(c)** Path straightness as gust amplitude changes, reusing the 100 intact neural states. Points and ribbons show means and pointwise 95% confidence intervals after averaging 20 gusts within each realization. Standard amplitude reproduces the original controller outcomes. This separate diagnostic contributes 10,000 controller runs, including 2,000 repeated standard-amplitude runs, and no new neural states. **(d)** Maximum absolute differences between intact and complete original reconstruction across the 100 paired realizations. All differences are zero. These are internal computational checks.

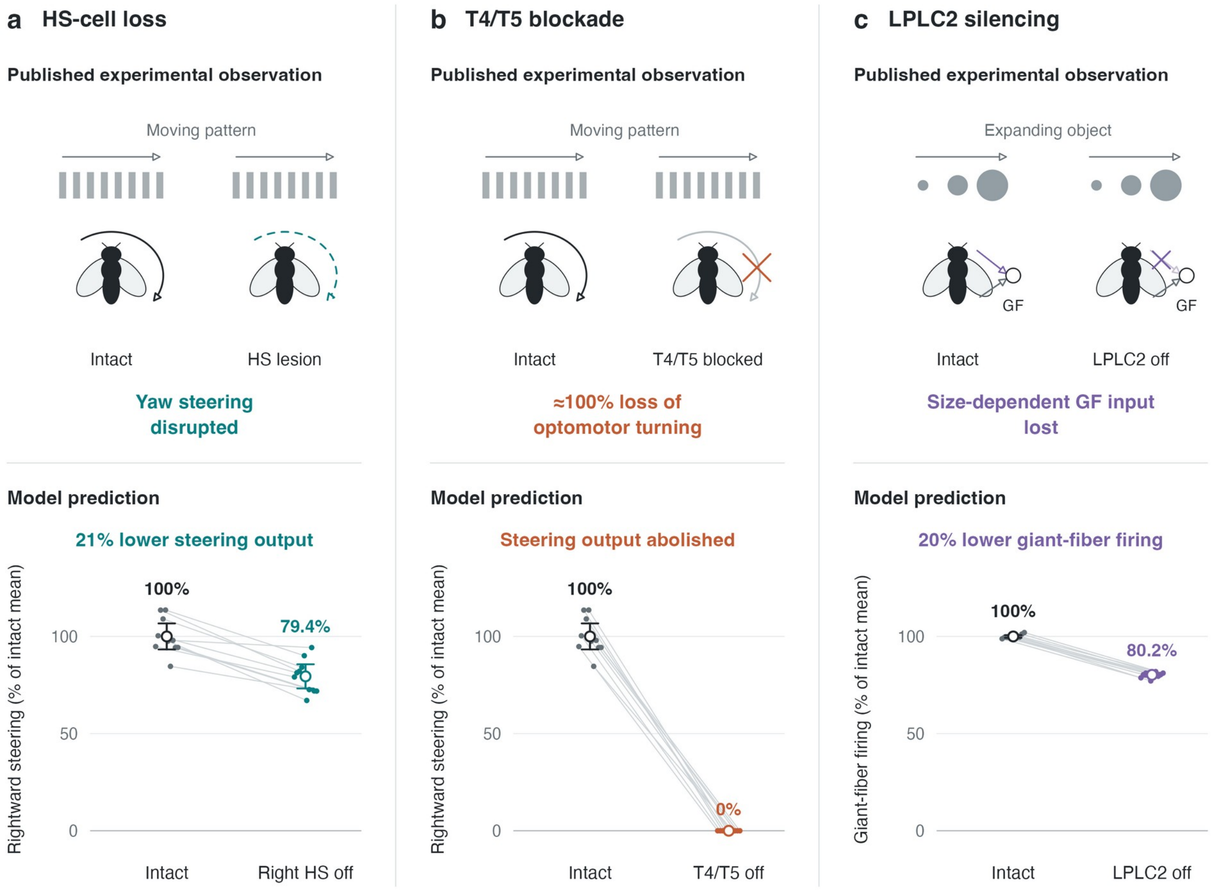


**Supplementary Figure 2: Selected experimental precedents for visual sensorimotor responses in the connectome model.** (a) Microsurgical HS lesions disrupted optomotor yaw steering in blowflies (ref. 20). Silencing the right HSN, HSE and HSS neurons in the Drosophila model reduced mean rightward steering output to 79.4% of the intact mean. (b) T4/T5 blockade abolished motion-following turning in walking Drosophila (ref. 21, Fig. 1b–d). The approximately 100% loss annotation summarizes the reported abolition of the optomotor response. Model T4/T5 silencing abolished steering output. (c) LPLC2 silencing removed the size-dependent component of the experimentally recorded giant-fiber (GF) looming response, while a velocity-dependent component remained (ref. 22). Purple and gray connections depict the lost size-related and retained inputs, respectively. Model LPLC2 silencing reduced mean GF firing from 249.0 to 199.8 Hz (80.2% retained) during combined LC4/LPLC2 drive.

Small points show all 10 sensory-event seeds in one connectome; lines connect matched intact and perturbed runs. Open circles and whiskers denote means and 95% t confidence intervals across seeds. Within each panel, every raw output is divided by the same observed intact mean, preserving intact-run variability; confidence intervals are scaled by that fixed denominator. Panels a and b show the raw rightward steering-decoder output; c shows GF firing. Parameters and decoder weights were unchanged. Fly icons are schematic. These are selected qualitative comparisons across different assays: T4/T5 are directly driven model inputs, and panel c tests pathway contribution without reproducing looming size/velocity dynamics or escape behavior.

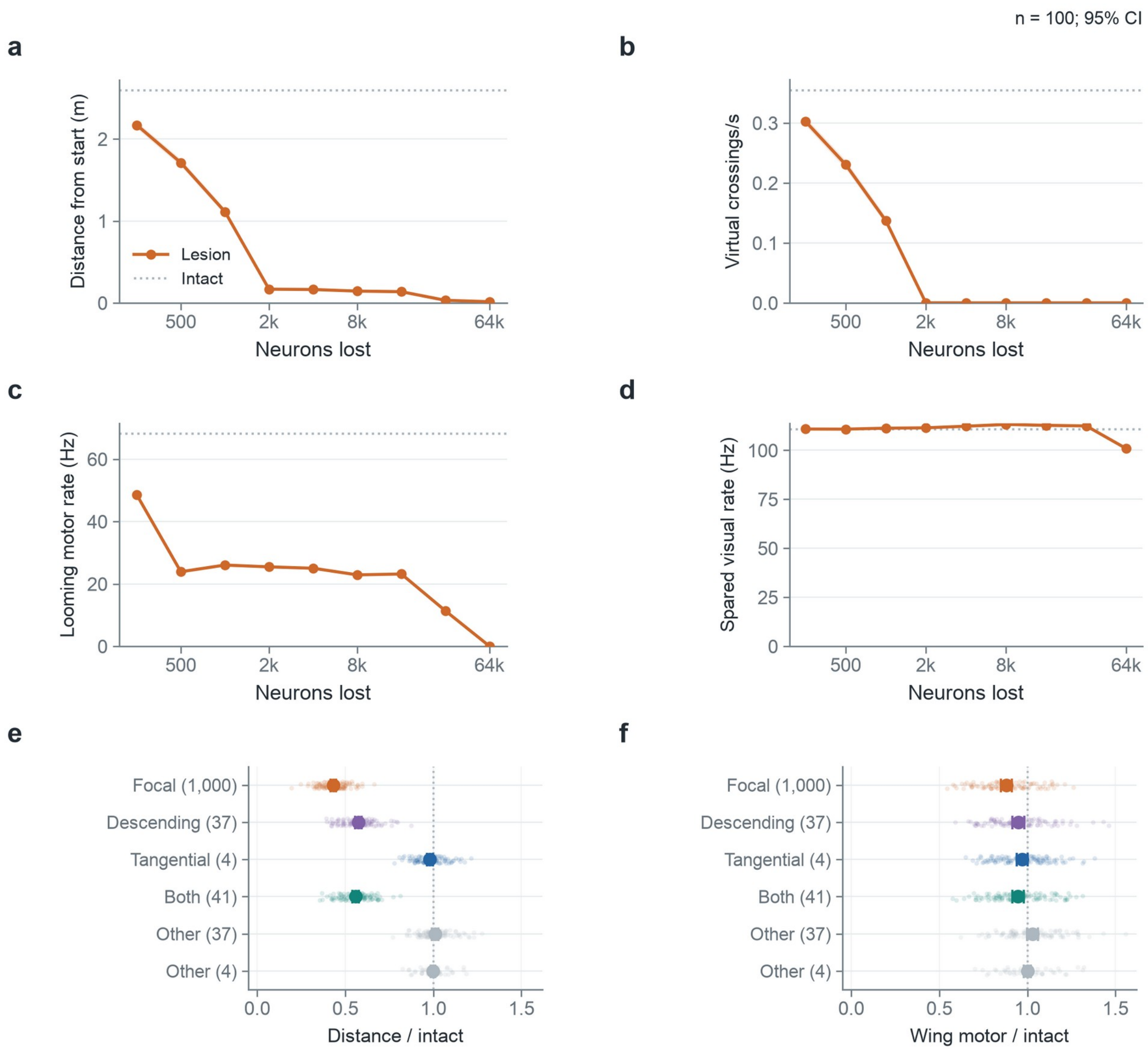


**Supplementary Figure 3: Absolute lesion readouts and cell-class contributions. (a–d)** Distance after ten seconds, repeated virtual crossings per second, looming-evoked motor rate and spared visual rate across the nested site-20 lesion series. Dotted gray lines show intact means. **(e,f)** Distance and wing-motor activity relative to matched intact after the 1,000-cell focal lesion or selective removal of its 37 descending neurons, four tangential cells, their 41-cell union, or 37 or four other cells from the same lesion mask. Dotted lines show intact. Points and uncertainty show means and pointwise 95% confidence intervals across 100 realizations; movement averages 20 matched gusts first. Cell-class ablations do not match focal-lesion geometry. Crossing rate counts repeated passages through virtual flight planes. Looming rates are neural activity.

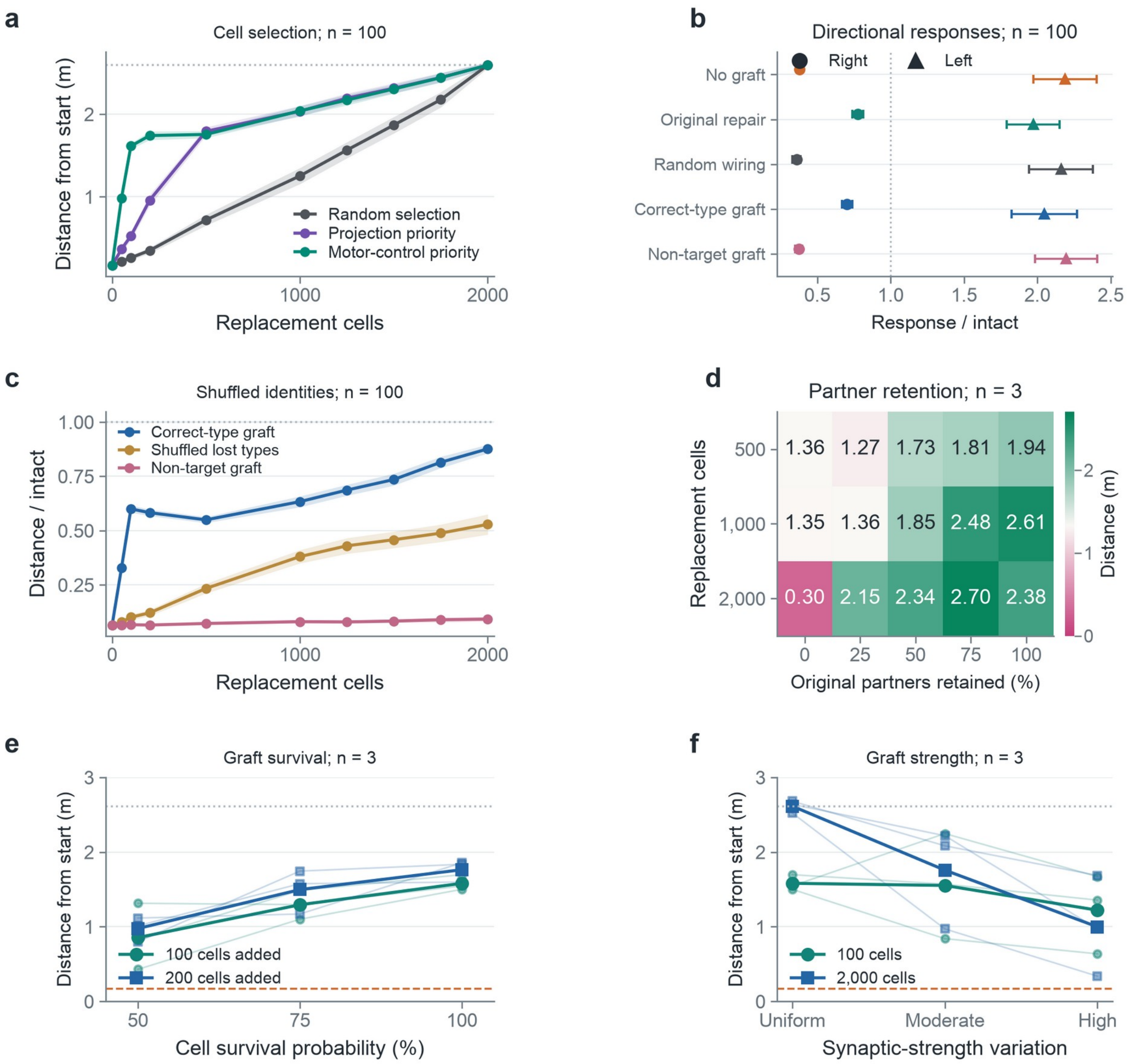


**Supplementary Figure 4: Additional replacement controls. (a)** Distance under random selection, projection priority or motor-control priority, all with original identities and available partners. Random selection differs from random wiring. **(b)** Rightward (circles) and leftward (triangles) responses for the five main repair conditions with 100 replacements, divided by each matched intact response. **(c)** Shuffled lost types, compared with correct-type and non-target grafts under the same donor-based wiring rule. Shuffling draws other types present in the lesion; non-target types are absent. Panels a–c use means and pointwise 95% confidence intervals across 100 realizations. **(d)** Earlier partner-retention grid: absolute distance after a 1,000-cell lesion, n = 3. The 2,000-cell graft duplicates templates and changes synaptic load. **(e)** Cell survival probability after adding 100 or 200 selected cells. **(f)** Uniform, moderately variable or highly variable synaptic strengths among 100 or 2,000 replacements, with all cells surviving. Lognormal multipliers have log standard deviations 0, 0.5 or 1 and expected multiplier one. Panels e,f show all three exploratory realizations and their means; faint lines join matched seeds. Gray dotted and orange dashed lines show intact and untreated means. All movement averages 20 gusts per realization.

### SUPPLEMENTARY MOVIES

**Supplementary Movie 1.** Lesion size and location. The first ten seconds compare 1,000-, 4,000- and 16,000-cell lesions at site 20 (top row) and site 25 (bottom row), with six matched 3D paths. The next 18 seconds reveal nine nested site-20 lesions, from 250 to 64,000 lost neurons, and all five **Fig. 1f** readouts. Each lesion size is a separate simulation. Points show means and ribbons the empirical central 95% simulation range across 100 realizations, with 20 gusts averaged for movement. Spared visual activity has a separate panel. The anatomy marks actual lesion masks.

**Supplementary Movie 2.** Neuronal responses to looming input, comparing intact, a 2,000-cell site-20 lesion and 100 replacements with original connections. Nine seconds of playback show 59–300 ms of the 300-ms trial at 30 fps. Purple marks input, blue host, teal replacement and orange motor activity, with an 8-ms display decay. Motor traces show the arithmetic mean of TTMn and DLMn population rates over the preceding 25 ms, matching the looming motor readout in the figures. Anatomy and rate axes share scale.

**Supplementary Movie 3.** Selecting cells to restore after a 2,000-cell site-20 lesion. Four matched ten-second paths compare no graft and 100 lost cells selected randomly, by projection priority or by motor-control priority. All replacements retain their original identities and available partners. Random selection chooses missing cells and differs from random wiring in **Fig. 2**. Colored anatomical points in the insets show the selected cells within the shared lesion. This comparison corresponds to **Supplementary Fig. 4a**.

**Supplementary Movie 4.** Replacement identity and wiring. Five **Fig. 2** conditions are compared: no graft, original repair, random wiring, correct-type graft and non-target graft. Separate ten-second chapters show 100 and 2,000 intended replacements after the same 2,000-cell lesion, matching **Fig. 2b**. Each dose is a separate simulation. Correct-type and non-target grafts copy partners from surviving cells of the assigned type using the same integration rule. Tables show mean distance, wing-motor and looming-motor output across 100 realizations, normalized to matched intact. Only complete original repair reproduces intact exactly; random wiring and non-target identity remain impaired at full dose. All panels share one camera and scale.
